# Artemether and Euphorbia Factor L9 suppress kynurenine production through distinct effects on Tryptophan metabolism

**DOI:** 10.1101/2025.06.02.657401

**Authors:** Alina L. Capatina, Tomasz Czechowski, Charlotte Plunkett-Jones, Thierry Tonon, Ioannis Kourtzelis, Benjamin R. Lichman, William J. Brackenbury, Ian A. Graham, Dimitris Lagos

## Abstract

L-Tryptophan (Trp) is an essential amino acid, catabolised through the kynurenine pathway, which is mediated by the enzymes indoleamine-2,3-dioxygenase 1 (IDO1), IDO2, or Trp-2,3-deoxygenase (TDO). Therapeutic targeting of Trp metabolism could be relevant to several pathologies. In cancer, IDO1 acts as an immune checkpoint suppressing effector T cell function. Yet, direct inhibition of IDO1 has had limited success in clinical trials. Therefore, alternative approaches to Trp metabolism therapeutic targeting are needed. We screened a library of 597 natural products (NPs) or NP derivatives for their effect on kynurenine production in triple negative breast cancer cells. This revealed 24 candidate inhibitors of kynurenine production. Amongst them, artemether, a member of the artemisinin family of anti-malarial drugs, suppressed kynurenine production, likely via an endoperoxide bridge-dependent mechanism. The Euphorbia factor L9 (EFL9) inhibited kynurenine production likely via a C7-benzoylation-dependent mechanism. Neither artemether nor EFL9 affected JAK/STAT signaling or IDO1 levels. Targeted metabolomics analyses demonstrated that artemether suppressed kynurenine production through heme sequestration, a mechanism that would affect all members of the IDO/TDO family of metalloenzymes. EFL9 affected purine and amino acid metabolism and the cellular redox balance. Comparisons to the effects of ouabain, a NP regulator of IDO1 levels, and Linrodostat, a clinically used small molecule IDO1 inhibitor, revealed distinct metabolic profiles, with ouabain and EFL9 showing the largest overlap. Importantly, the kynurenine-suppressing activity of artemether and EFL9 is not cancer cell-specific. Overall, our findings set the foundation for the use of derivatives of artemether or EFL9 as novel Trp metabolism-targeting therapeutics.

## Introduction

The catabolism of L-Tryptophan (Trp) by antigen presenting, infected, or cancer cells represents a well-studied metabolic immune checkpoint pathway with therapeutic potential (1–3). Trp is an essential amino acid with low abundance in the body (4, 5). It is assimilated from nutrients and can serve a range of physiological functions, including protein synthesis and production of neuroactive metabolites required for serotonin synthesis (1). Over 90% of available Trp is catabolised through the kynurenine pathway (2). The first step of the kynurenine pathway is mediated by the enzyme indoleamine-2,3-dioxygenase (IDO). Depending on the tissue, one of two isoforms can be expressed, IDO1 or IDO2. In the liver, the first step of the kynurenine pathway and rate-limiting reaction is performed by Trp deoxygenase or TDO. However, IDO1 is also present in the liver and seems to be exacerbated in disease, playing an essential role in regulating the local immune landscape (6). Downstream metabolites of the pathway are involved in energy metabolism (*de novo* NAD^+^), neuromodulation (e.g. kynurenic acid, quinolinic acid), or immuno-regulation (e.g. kynurenic acid, kynurenine) (7). Inhibiting Trp metabolism has been identified as a potential immune checkpoint-targeting therapy. By depleting Trp, IDO1/2 and/ or TDO are deprived of their primary substrate and thus kynurenine production is impaired, preventing the immunosuppressive effects of kynurenine, including inhibition of effector T cells through Trp and glucose starvation, and preferential differentiation of regulatory T cells (8–11). Metabolites downstream of kynurenine, such as kynurenic acid, xanthurenic acid and cinnabarinic acid interact with the aryl hydrocarbon receptor (AhR). Furthermore, Trp depletion has been shown to contribute to neoantigen formation by interfering with translation (12). The same translational defects can also trigger activation of general control non-derepressible-2 (GCN2) dependent starvation responses, which impair growth and activate autophagy pathway, likely to clear cancerous mutations and thus decrease tumour progression (1, 2, 13).

In breast cancer, the rate-limiting and Trp catabolic enzyme IDO1 is highly upregulated (14, 15). In fact, IDO1 is significantly upregulated in most solid cancers, functioning as a prognostic biomarker of disease progression (16, 17). Mechanistically, IDO1 expression in cancer cells is induced by proinflammatory stimulation via the IFNγ receptor (IFNγR). This activates the JAK1/2-STAT1 pathway which is required for IDO1 expression. TNF further amplifies IDO1 levels (17–20). IDO1 converts Trp into an alkyl-phenylketone called N-formylkynurenine, further converted by formamidase to kynurenine. In this way, IDO1 induces Trp starvation of cytotoxic T cells, preventing attack against the tumour, as well as promoting differentiation of regulatory T cells via kynurenine-AhR regulated transcriptional reprogramming (21). Energetically, the kynurenine pathway ultimately generates *de novo* NAD^+^ molecules, thus contributing to growth and survival of IDO1-expressing cells (22). For these reasons, a series of small molecule inhibitors that interact directly with IDO1 such as Indoximod, Epacadostat, Linrodostat mesylate, Navoximod or PF-0684003, as well as peptide vaccines, have been designed for IDO1-targeted therapies (23, 24). Of these, Indoximod, a Trp mimetic, has been clinically approved for the treatment of melanoma in combination with chemotherapy (23). The limited clinical efficacy through direct IDO1 inhibition supports the need for novel approaches of targeting the kynurenine response, such as interfering with Trp metabolism, or targeting other linked physiological parameters, such as ionic dynamics (25).

Natural products (NPs) have been selected through millions of years of evolution to be bioactive. They account for almost a quarter of all drugs (50% in cancer) in terms of origin or inspiration (26). NPs have high chemical and stereochemical flexibility and have been used for cancer treatment, examples include the topoisomerase I inhibitor camptothecin, currently FDA approved for the treatment of breast, ovarian, lung and colon cancers, or taxol derivatives which work as microtubule inhibitors (27). Interestingly, some natural compounds have also been identified as potential IDO1 inhibitors with mild/ moderate activity: natural quinones, such as ubiquinone, which has been synthetically modified to improve its potency, or vitamin K3, also known for its function in regulating cell growth (28–30). Flavonoids reduced survival of human cervical cancer cells by direct interaction with IDO1 (31), and curcumin has been shown to supress JAK/STAT-mediated IDO1 expression (32, 33).

Breast cancer shows limited responses to immune checkpoint inhibitors (34–36), with most common treatment options being chemotherapy, surgery, hormone therapy as well as some targeted therapies such as Human epidermal growth factor receptor 2 (HER2) and poly-ADP ribose polymerase (PARP) inhibitors. However, the most severe type of breast cancer, triple negative breast cancer (TNBC), has been shown to have increased immune infiltration and thus better response to immunotherapy, leading to FDA approval of the anti-PD-1 monoclonal antibody therapy, Pembrolizumab, for early-stage TNBC treatment (37–39). The limited response of TNBC to conventional treatments highlights the need for developing better immune checkpoint targeted therapies for these patients. We have previously characterised the inhibitory effect on IDO1 in TNBC cell lines of two natural compounds, digoxin, first identified in foxglove (*Digitalis lanata*), and a compound extracted from the *Acokanthera schimperi* and *Strophanthus gratus* plants, ouabain, both members of the cardiac glycoside family. We showed that cardiac glycosides are indirect modulators of kynurenine production, causing decreased activation of the main IDO1 transcriptional regulator, STAT1, potentially achieved via dysregulation of the intracellular Na^+^ concentration (25). Here, using an assay that quantifies kynurenine secretion, we screened a library of 597 NPs or NP derivatives. We identified and validated two compounds that inhibited kynurenine production and Trp metabolism: artemether, an acetal derivative of the endoperoxide-containing sesquiterpene artemisinin, and Euphorbia factor L9 (EFL9). Mechanistically, artemether, but not EFL9, inhibited kynurenine production through a heme-dependent mechanism, a strategy that could potentially affect all metalloenzymes of the IDO/TDO family.

## Results

### A screen of 597 unique NPs or derivatives identifies 24 inhibitors and 1 enhancer of kynurenine production in TNBC cells

We have previously identified the cardiac glycoside drugs, ouabain and digoxin, as inhibitors of the kynurenine response (25) using a colorimetric assay (40, 41). Using the same assay, we screened the effect of a library of 597 NPs or NP derivatives on kynurenine production by MDA-MB-231 TNBC cells following treatment with IFNγ and TNF. The combined compound library consisted predominantly of terpenoids with other classes of compounds such as: alkaloids, phenylpropanoids, polysaccharides and casbene-derivatives (**Supplementary Figure S1**). The full list of compounds is available in **Supplementary Table S1A**. The first round of screening included all compounds and was used to identify compounds of interest (COIs), which affected kynurenine production without impacting on cell viability. Ouabain was used as a NP positive control given its effect on kynurenine production (25). We identified COIs either based on fold-change in kynurenine levels (2-fold decrease or 1.4-fold increase) compared to the Dimethyl sulfoxide (DMSO)-treated cells (ran in triplicate in every plate) or based on a change in kynurenine levels by more than three standard deviations (SD) from the mean of the DMSO-treated control. These arbitrary thresholds were used to allow selection of a range of compounds for further validation. With regards to fold-change, we initially aimed to select kynurenine fold-change thresholds equivalent to 50% change either side of the control. However, due to the low resolving power of the assay at high kynurenine concentrations, we selected a 40% increase as an initial threshold for detecting potential enhancers of kynurenine production. With regards to SD difference, only plates that had a 3xSD difference between the positive (ouabain) and negative (DMSO) control were included in the analysis (see Methods) (42). Each compound was tested in two independent replicates, and the workflow of the pharmacological screen is shown in **Figure 1A**. We identified 50 COIs (shown in red in **Figure 1B**), 44 of which caused a reduction in kynurenine production. All inhibitory and enhancer COIs gave a viability value equal to or over 0.7 of that corresponding to the vehicle unstimulated control (**Figure 1C**). The kynurenine, fold change, SD variation and viability data for all tested compounds included in the screen, as well as for the controls, are shown in **Supplementary Table S1A and B**.

**Figure 1:**
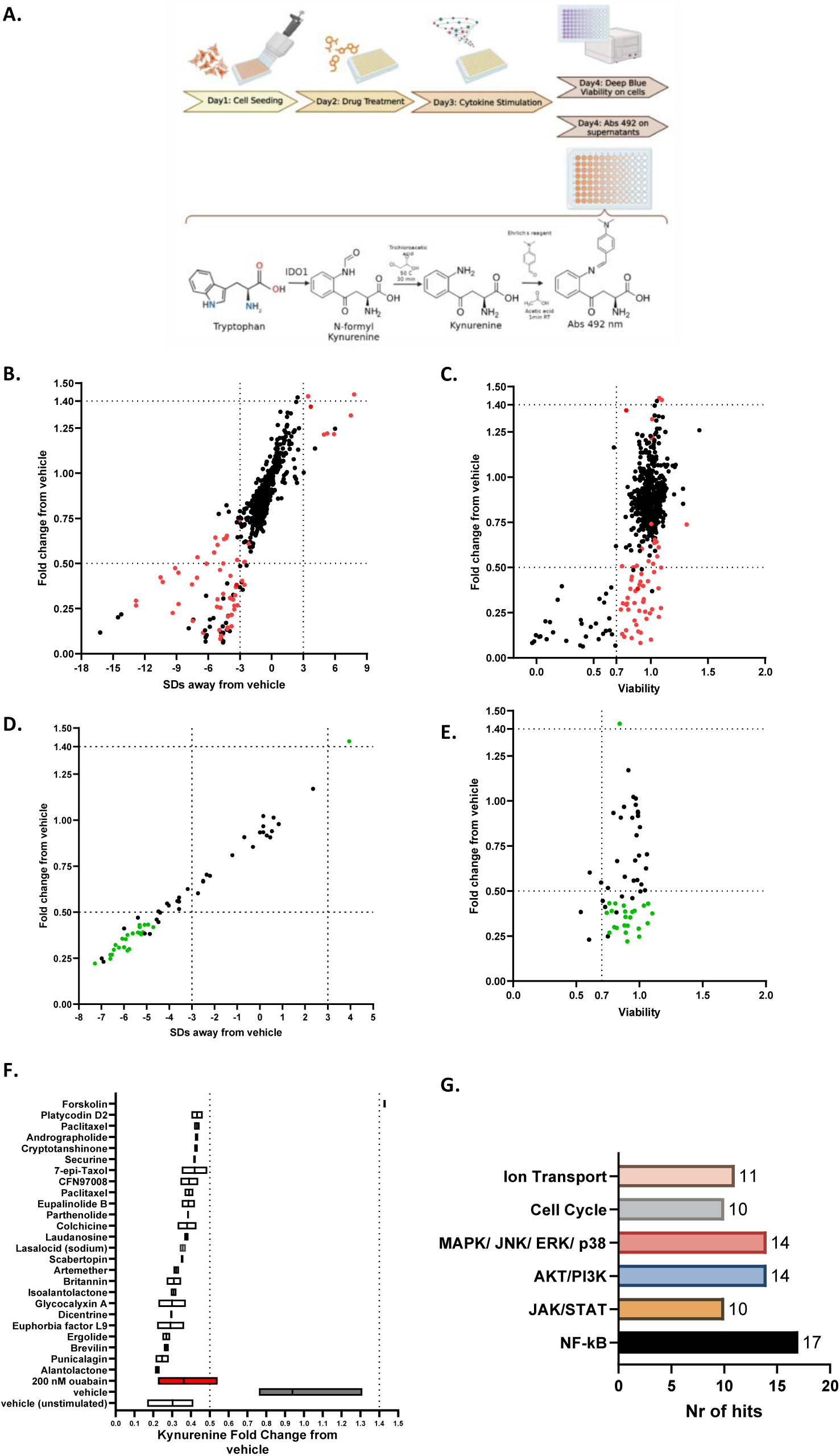
Screening a NP library identifies candidate regulators of kynurenine production in MDA-MB-231 cells. **A.** Drug screen workflow, including the kynurenine and viability assays. **B.** First screening round: Fold change in kynurenine levels normalised to the DMSO-treated cells control vs variation from mean levels of DMSO-treated cells measured in SDs. Average values for fold change and SD variation from the kynurenine levels in the DMSO-treated cells were plotted (n=3 for DMSO-treated cells per plate and n=2 for all other tested compounds). **C.** Fold change in kynurenine levels normalised to the vehicle stimulated positive control vs viability (n=2). **D.** Validation round: Fold change in kynurenine levels normalised to the DMSO-treated cells vs variation from mean levels of DMSO-treated cells measured in SDs from the mean of DMSO-treated cells (n=3 for DMSO-treated cells and n=2 for all other tested compounds). **E.** Fold change in kynurenine levels normalised to the vehicle stimulated positive control vs viability (n=2 biological replicates, plotted as averages). **F.** List of hits ordered by impact on kynurenine fold change, data were plotted as mean and range (n=2). **G.** Natural compound screen hits and their previously published biological activities – Representation of the main pathways affected by the hits, and number of compounds reported to affect each pathway. For **B** and **C**: red indicates compounds selected for further validation (COIs and a selection of compounds with borderline effects on kynurenine production). For **D** and **E** green indicates validated compounds (hits).

All the COIs and an arbitrarily selected set of 8 borderline compounds (for example, compounds that resulted in a 0.51 decrease in kynurenine, or viability of 0.69) were chosen for further validation (58 compounds in total). For the validation round, we designated as inhibitory hit compounds those that caused a 2-fold decrease in kynurenine reduction and a decrease greater than mean-3xSD of the DMSO control. As enhancers, we designated compounds that increased kynurenine production by 1.4-fold and mean+3xSD of the DMSO-treated control. This identified 24 inhibitory hits, and 1 enhancer (Forskolin) hit (shown in green in **Figure 1D**). All hits had a viability score above the 0.7 threshold (**Figure 1E**). **Supplementary Table S1A** shows the data for each compound tested in the validation round. The identified hits were ordered based on the magnitude of the kynurenine production inhibition (**Figure 1F**) and a literature search identified a list of signalling pathways previously linked to these compounds. These included NF-kB signalling, JAK/STAT activation, AKT/PI3K signalling, MAPK/JNK/ERK/p38 activation, cell cycle regulation and ion transport mechanisms (**Figure 1G** and **Supplementary Table S2**).

### Artemisinin derivatives that carry an endoperoxide structure suppress kynurenine production without impacting on expression of IDO1 and the JAK/STAT pathway

Having identified 24 compounds with inhibitory activity against the kynurenine response, the next step was to select candidates for further mechanistic studies. We selected compound classes that exhibited a correlation between their chemical structure and their biological activity, indicating a potential role of specific chemical groups in modulating the kynurenine response. For each compound class, the most potent kynurenine inhibitory compounds were selected for further studies, together with a structurally similar but biologically inactive member of the class, selected as a negative control. The first class of compounds that we focused on consisted of artemisinin derivatives, traditionally used as anti-malaria therapeutics (43, 44).

All compounds that carried the oxygen endoperoxide bridge (**Figure 2A**), highlighted with the red box, demonstrated activity either as COIs (artemether, artemisinin, artesunate and dihydro-artemisinin) or as validated hits (artemether). The endoperoxide bridge has been identified as the moiety responsible for the anti-malaria activity in artemisinin derivative molecules (44, 45). Notably, deoxyartemisinin, an artemisinin derivative that lacks the endoperoxide core (**Figure 2A**), was also included in the screen and showed no activity against the kynurenine response. Titration of artemether and deoxyartemisinin showed that only the former induces a concentration-dependent decrease in kynurenine production (**Figure 2B**), without affecting viability. Artemether had no impact on IDO1 mRNA or protein levels (**Figure 2C-F**). Consistent with this finding, artemether did not affect activation of the JAK/STAT pathway or expression of PD-L1, another immune checkpoint protein downstream of the same pathway (**Figure 2D**). This suggests that, unlike our previous findings with ouabain (25), artemether does not reduce kynurenine by interfering with IFNγ signalling and subsequent IDO1 expression. Overall, artemether suppressed kynurenine production correlating with a potential activity from the endoperoxide bridge structure within the molecule of artemether, without affecting JAK/STAT signalling or IDO1 levels.

**Figure 2:**
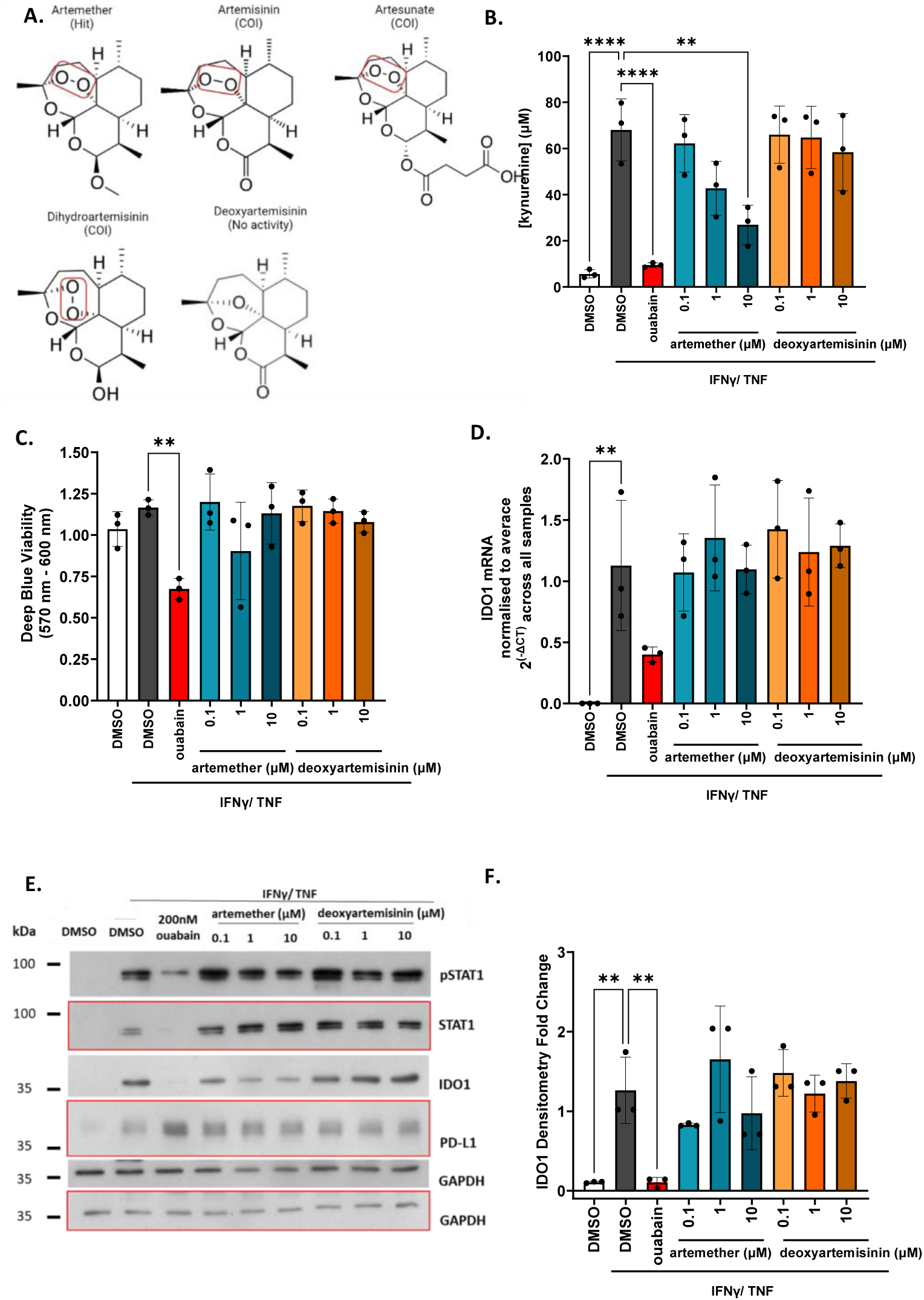
Endoperoxide bridge containing artemisinin derivatives suppress kynurenine production in TNBC cells without affecting IDO1 expression. **A.** Chemical structures of artemisinin derivatives included in the screen. **B.** Effect of artemether and deoxy artemisinin titration on the kynurenine levels in MDA-MB-231 cells (n=3 biological replicates). **C.** As in B but for viability (n=3 biological replicates). **D.** As in B but for IDO1 mRNA levels, samples were normalized to β-actin (n=3 biological replicates). **E.** Western blot showing the effect of artemether and deoxyartemisinin on (p)STAT1, IDO1 and PD-L1; bands corresponding to the same blot were marked with red margins or no margins (representative for n=3 biological replicates). **F.** Densitometry fold change for IDO1 protein levels (n=3 biological replicates). All experiments presented in this figure used ouabain at 200 nM as a positive control. All data were plotted as mean and SD, statistical comparisons were carried out using a one-way ANOVA coupled with a Bonferroni’s multiple comparison test (all conditions were compared to the DMSO IFNγ/TNF) (**p<0.01; ****p<0.0001). Legend: COI – compound of interest. US – unstimulated.

### Euphorbia factors with a C7 modification suppress kynurenine production without impacting on expression of IDO1 and the JAK/STAT pathway

We examined a series of diterpenoid NPs from *Euphorbia* sp. with a lathyrane skeleton (**Figure 3A**). We identified EFL2 as a COI, with some activity on the kynurenine response, while EFL9 was identified as a hit (**Supplementary Table S1A**). EFL1 and all the other members of the class included in the screen had no impact on kynurenine. EFL9 has not been widely researched, and little is known about its biological activity. Structurally, EFL2 and EFL9 share a C7 modification, which is not found in any of the other class members included in this screen (**Figure 3A**), suggesting that the C7 modification might be responsible for their effect on kynurenine production. Titration of the hit (EFL9), the COI (EFL2), and another member of the class with similar structure but no C7 modification (EFL3) was carried out. **Figure 3B** shows that both EFL9 and EFL2 suppress kynurenine production in a dose-dependent manner, with EFL9 being more potent. None of the three factors tested affected viability (**Figure 3C**), IDO1 mRNA levels (**Figure 3D**), or IDO1, (p)STAT1 and PD-L1 protein levels (**Figure 3E, F**). This result demonstrated that EFL2 and EFL9 inhibit kynurenine production likely via a C7-benzoylation-dependent mechanism.

**Figure 3:**
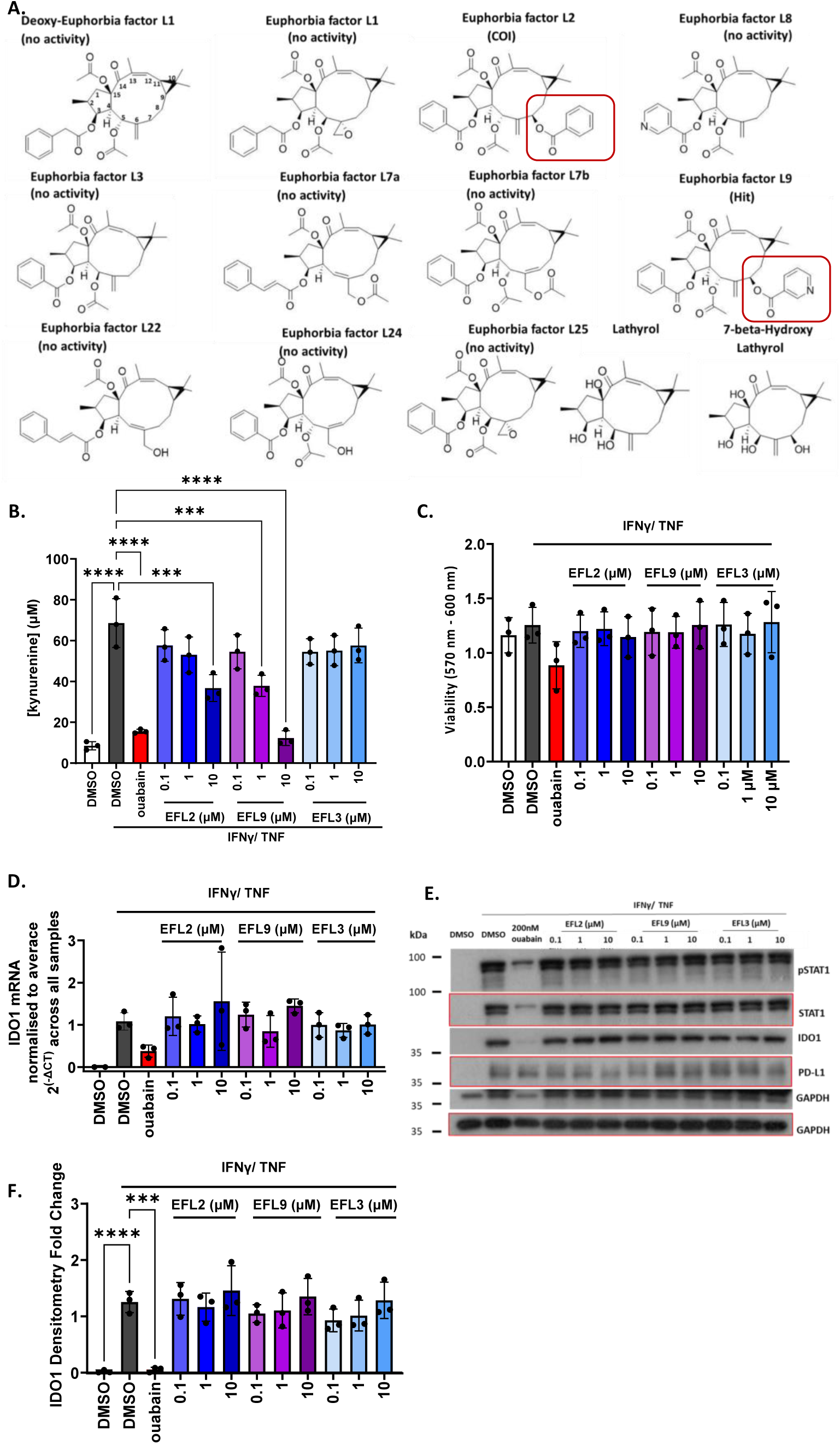
Euphorbia factors L2 and L9 suppress kynurenine production from MDA-MB-231 cells, without affecting IDO1 expression. **A.** Chemical structures of Euphorbia factors included in the screen. C7 modifications on EFL2 and EFL9 are shown. **B.** Effect of EFL9, EFL2 and EFL3 titration on the kynurenine response in MDA-MB-231 cells (n=3 biological replicates). **C.** As in B but for viability (n=3 biological replicates). **D.** As in B but for IDO1 mRNA levels. Levels were normalized to β-actin and average expression value across all the samples (n=3 biological replicates). **E.** Western blot showing the effect of EFL9, EFL2 and EFL3 on (p)STAT1, IDO1 and PD-L1; bands corresponding to the same blot were marked with red margins or no margins (representative for n=3 biological replicates). **F.** Densitometry fold change for IDO1 protein levels (n=3 biological replicates). All experiments presented in this figure used ouabain at 200 nM as a positive control. All data were plotted as mean and SD and statistical comparisons were carried out on B, C and F using a one-way ANOVA coupled with a Bonferroni’s multiple comparison test (all conditions were compared to the DMSO IFNγ/TNF) (***p<0.001; ****p<0.0001).

### Artemether and EFL9 and EFL2 suppress kynurenine production from non-transformed primary mammary epithelial cells

Having shown that artemether and EFL9 and EFL2 decrease kynurenine levels in MDA-MB-231 cells, we tested whether these effects were restricted to cancer cells. The effect of ouabain, artemether (and its control deoxyartemisinin), EFL9 and EFL2 (and their control EFL3) on kynurenine and IDO1 expression were measured in primary human mammary epithelial cells (HMEC) from two different donors. We found that kynurenine levels in HMECs follow the same trend as in MDA-MB-231 cells in response to artemether, EFL9, and ouabain (**Figure 4A**). None of the compounds affected cell viability (**Figure 4B**). As in the case of breast cancer cells, ouabain caused a decrease in IDO1 mRNA and protein levels, whereas artemisinin derivatives and Euphorbia factors did not affect IDO1 expression (**Figure 4C, D**). These results demonstrated that the effects of ouabain, artemether, EFL9 and EFL2 on kynurenine are not specific to cancer cells, suggesting that these compounds might affect core processes involved in Trp metabolism and kynurenine production.

**Figure 4:**
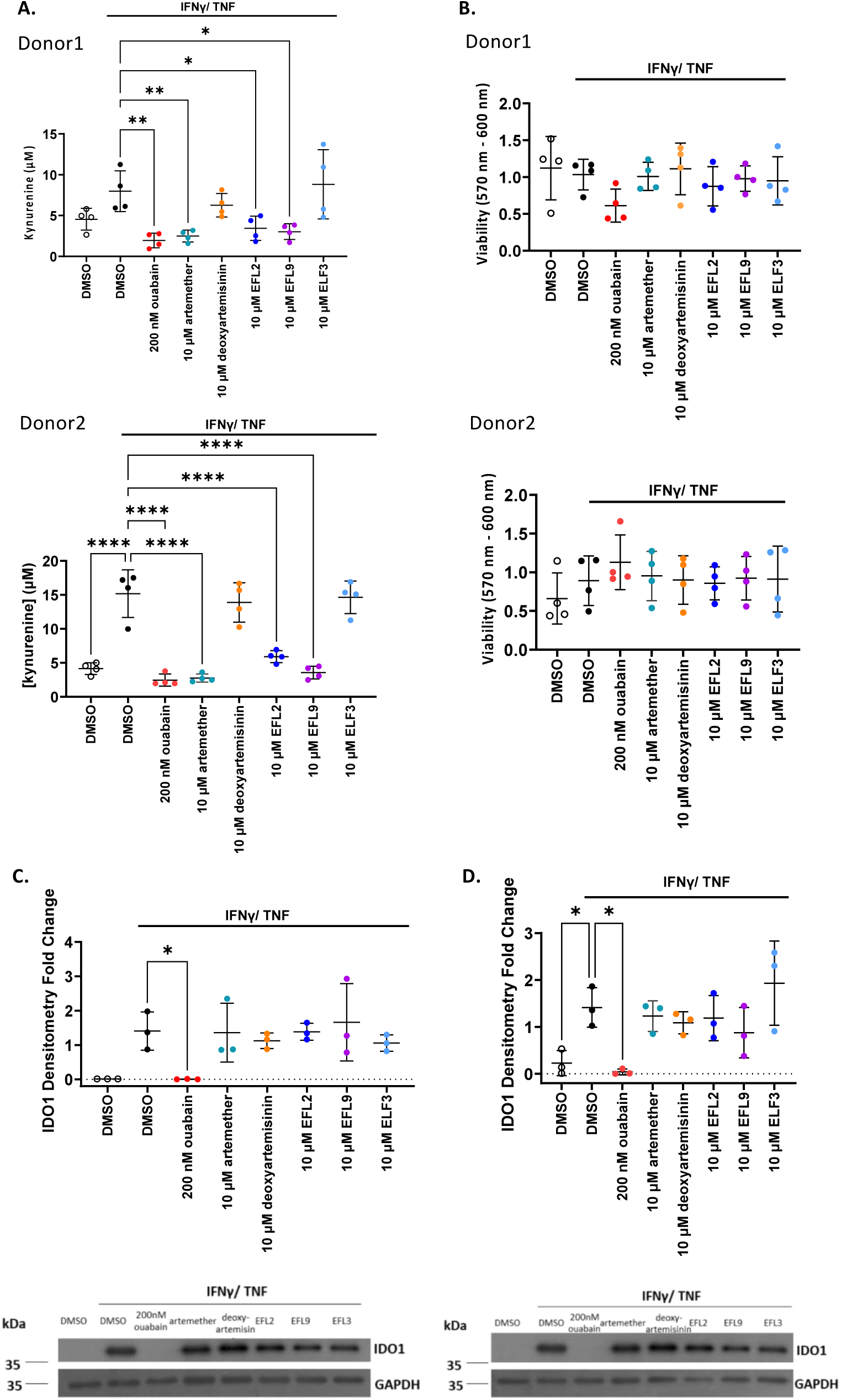
Ouabain, artemether, EFL9 and EFL2 suppress kynurenine production and IDO1 expression in primary HMECs. **A.** Effect of ouabain, artemisinin-derivatives and EFL9, EFL2 and EFL3 on the kynurenine response in HMECs (n=4 experimental replicates for each donor). **B.** As in A but for viability (n=4). **C.** As in A for IDO1 protein levels and densitometry analysis in donor 1, one representative western blot as included (n=3). **D.** As in A for IDO1 protein levels and densitometry analysis in donor 2, one representative western blot as included (n=3). All data were plotted as mean and SD, statistical comparisons were carried out using a one-way ANOVA coupled with a Bonferroni’s multiple comparison test (all conditions were compared to the DMSO IFNγ/TNF) (*p<0.05, **p<0.01; ****p<0.0001).

### Artemether, EFL9, and ouabain have overlapping and distinct effects on Trp metabolism in TNBC cells

Having shown that artemether and EFL9 affect kynurenine through IDO1 expression-independent mechanisms, we further assessed their impact on Trp metabolism. To this end, we measured intracellular levels of Trp metabolites by mass spectrometry in control (DMSO-treated) cells and in response to treatment with artemether and EFL9. EFL9 was selected for these experiments as it showed the most potent effect on kynurenine (**Figure 3B**). We also compared the metabolic profiles of artemether- and EFL9-treated cells to those of cells treated with ouabain or the clinically used small molecule IDO1 inhibitor, Linrodostat (46–48).

Principal component analysis (PCA) was carried out on all the detected metabolites (135 metabolites) and showed a clear separation of the 4 treatment conditions, with some similarity observed between the artemether and DMSO conditions (**Supplementary Figure S2**). The data set was then reduced to metabolites that could be confidently annotated (47 metabolites).

There was still separation between the four groups (**Figure 5A**), albeit weaker likely due to reducing the number of included metabolites. The EFL9 group seemed to be the most heterogenous, as well as the most distinct from the DMSO group, while artemether-treated cells showed the highest similarity to the DMSO condition. Finally, ouabain treated samples gave the tightest grouping based on PC1, showing some overlap with the EFL9 samples (**Figure 5A**).

**Figure 5:**
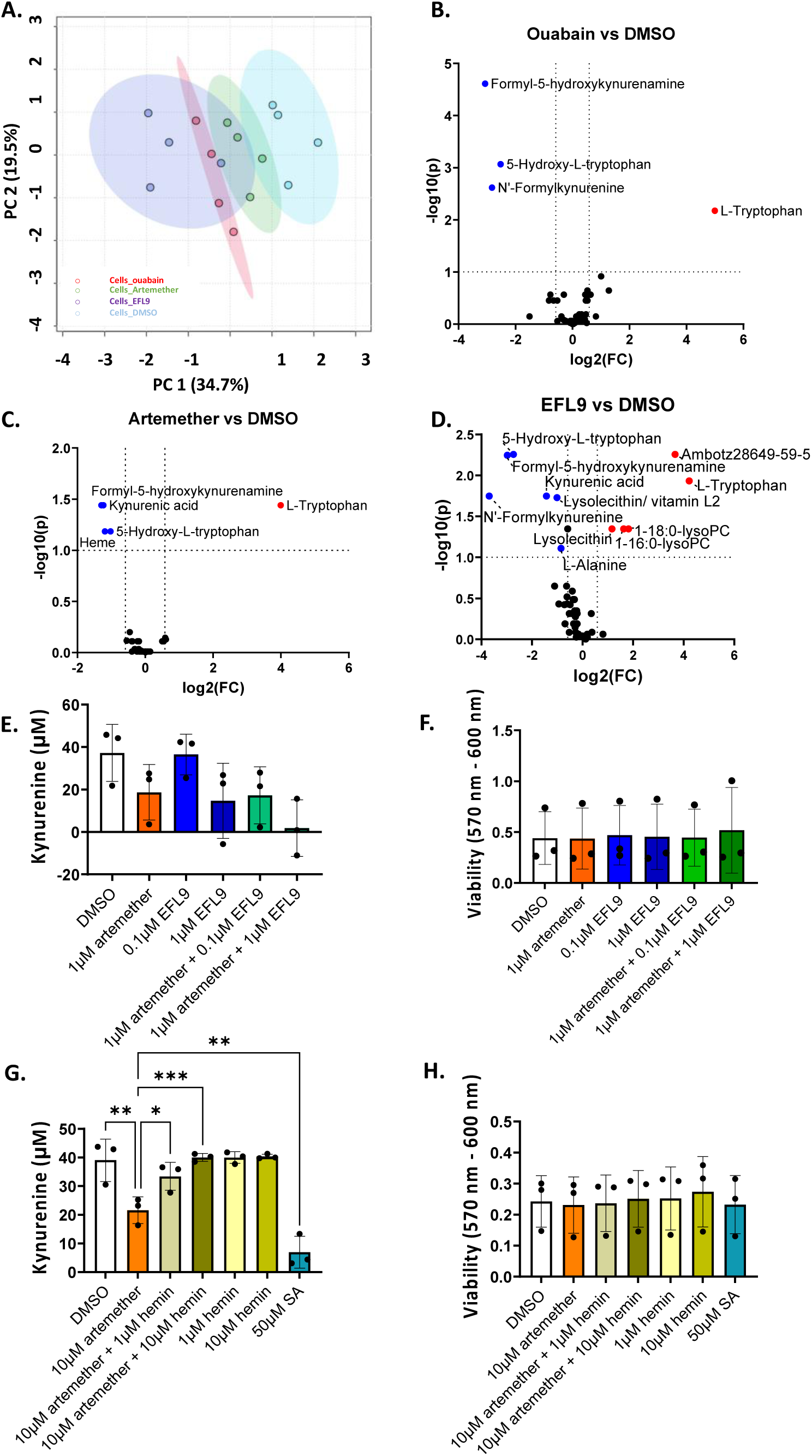
Impact of ouabain, artemether and EFL9 on Trp metabolites in MDA-MB-231 cell extracts. **A.** PCA analysis of the annotated metabolite data set (47 metabolites), n=4 biological replicates. **B.** Volcano plots of log_10_Fold change vs log_10_(False Discovery Rate, FDR) value for metabolites in response to ouabain, indicating statistically affected metabolites, n=4 biological replicates. **C.** As in B for artemether. **D.** As in B for EFL9. Analysis was carried out using the Metaboanalyst software. **E.** Kynurenine levels in response to hemin in DMSO- or artemether-treated MDA-MB-231 cells, and in response to succinyl acetone (SA; n=3). **F.** As in E but for viability (n=3). For A to D, data were median normalized and log_10_ transformed, with no further filtering applied. For volcano plots, the fold change threshold was set at 1.5, while the FDR threshold was set at 0.1. For E and F, mean and SD is shown. Statistical analysis was carried out using a one-way ANOVA coupled with a Bonferroni’s post-test (E, comparing specific pairs of data) and a Tukey’s post-test (F) (*p<0.05, **p<0.01; ****p<0.0001).

Next, we performed pair-wise comparisons between the cell lysate metabolites in the DMSO-treated group and the three drug-treated groups. These analyses included the annotated metabolites only. Ouabain treatment led to statistically significant changes in the levels of 4 metabolites (**Figure 5B**), whereas artemether and EFL9 treatment resulted in statistically significant changes in 5 and 11 metabolites, respectively (**Figure 5C, D**). All drugs led to an increase in the Trp levels in the cell, while decreasing the levels of formyl-5- hydroxykynurenamine and 5-hydroxy-L-Trp. Ouabain and EFL9 decreased the levels of N-formylkynurenine. Both artemether and EFL9 also suppressed levels of kynurenic acid. Interestingly, only artemether decreased intracellular heme levels. EFL9 also decreased the levels of 5’-methylthioadenosine and L-alanine.

Having observed overlaps between the effects of artemether and EFL9 on Trp metabolism in MBA-MB-231 cells, we explored their combinatory effect. We found that artemether and EFL9 have additive effects on kynurenine suppression (**Figure 5E, F**), suggesting that they operate through independent mechanisms. Our metabolic profiling experiment had indicated that only artemether treatment led to a decrease in heme levels. Indeed, restoring heme levels through addition of hemin reversed the effect of artemether without affecting kynurenine levels in control cells (**Figure 5G, H**). In agreement with this, inhibition of heme biosynthesis with succinyl acetone (SA) also led to a reduction in kynurenine levels in MDA-MB-231 cells (**Figure 5G, H**). We also measured levels of secreted metabolites in culture supernatants in response to ouabain, artemether and EFL9 (**Supplementary Figure S3**). All three drugs showed a decrease in N-formylkynurenine, formyl-5-hydroxykynurenamin and 5-hydroxy-L-tryptophan, while the levels of Trp in the supernatants were elevated in comparison to the DMSO control (**Supplementary Figures S3 and S4**). This indicated that the intracellular reduction in N-formylkynurenine was not due to increased export of kynurenine but a result of kynurenine pathway inhibition. Taken together, these results reveal that artemether and EFL9 have overlapping and distinct effects on Trp metabolism and kynurenine production. Artemether controls kynurenine production through heme sequestration whereas EFL9 causes a high number of metabolic changes, affecting purine and amino acid metabolism in the cell.

### Comparison of changes in Trp metabolism due to NP kynurenine production inhibitors and the direct IDO1 inhibitor Linrodostat

Our analyses gave us the opportunity to compare the metabolic effects of NP and NP-derived inhibitors of kynurenine production (artemether, EFL9, and ouabain) and of Linrodostat in MDA-MB-231 cells. We confirmed that Linrodostat suppressed kynurenine levels in a concentration dependent manner, without impacting on viability (**Supplementary Figure S5A**). Metabolomics analysis on cell lysate samples treated with Linrodostat (155 metabolites detected, 28 of those being confidently annotated) demonstrated clear clustering of control and Linrodostat-treated cells (**Supplementary Figure S5B**, for annotated metabolites). Differential abundance analysis showed that Linrodostat treatment caused statistically significant changes in 16 metabolites (**Supplementary Figure S5C**).

Integrating our metabolomics analyses, we mapped the changes in intracellular metabolites occurring due to ouabain (**Figure 6A**), Linrodostat (**Figure 6B**), artemether and EFL9 (**Figure 6C**). Of note, ouabain, which suppresses IDO1 levels, and Linrodostat, which inhibits IDO1 activity, demonstrated completely overlapping profiles in terms of downregulated metabolites. Ouabain downregulated the levels of the immediate IDO1 reaction product, N-formylkynurenine, as expected based on its negative effect on IDO1 expression (25). Ouabain also impacted on the serotonin/indole pathway, reducing metabolites such as formyl-5- hydroxykynurenamine, which is a serotonin-derived IDO-enzyme reaction product, but also upstream metabolites such as 5-hydroxy-L-Trp, which is a direct metabolite of Trp produced by Trp hydroxylases 1 and 2 (TPH1/2). Alongside Trp, Linrodostat also upregulated a unique range of key Trp metabolites, including formylanthranilic acid and indole derivatives such as 5-hydroxyindoleacetaldehyde/indole acetic acid and indoleacetaldehyde. In addition, Linrodostat also upregulated other metabolites such as vitamin B6 (pyridoxine/ adermine), purine/energy carrying metabolites (adenosine monophosphate, isonicotinamide) and amino acid levels (valinium).

**Figure 6:**
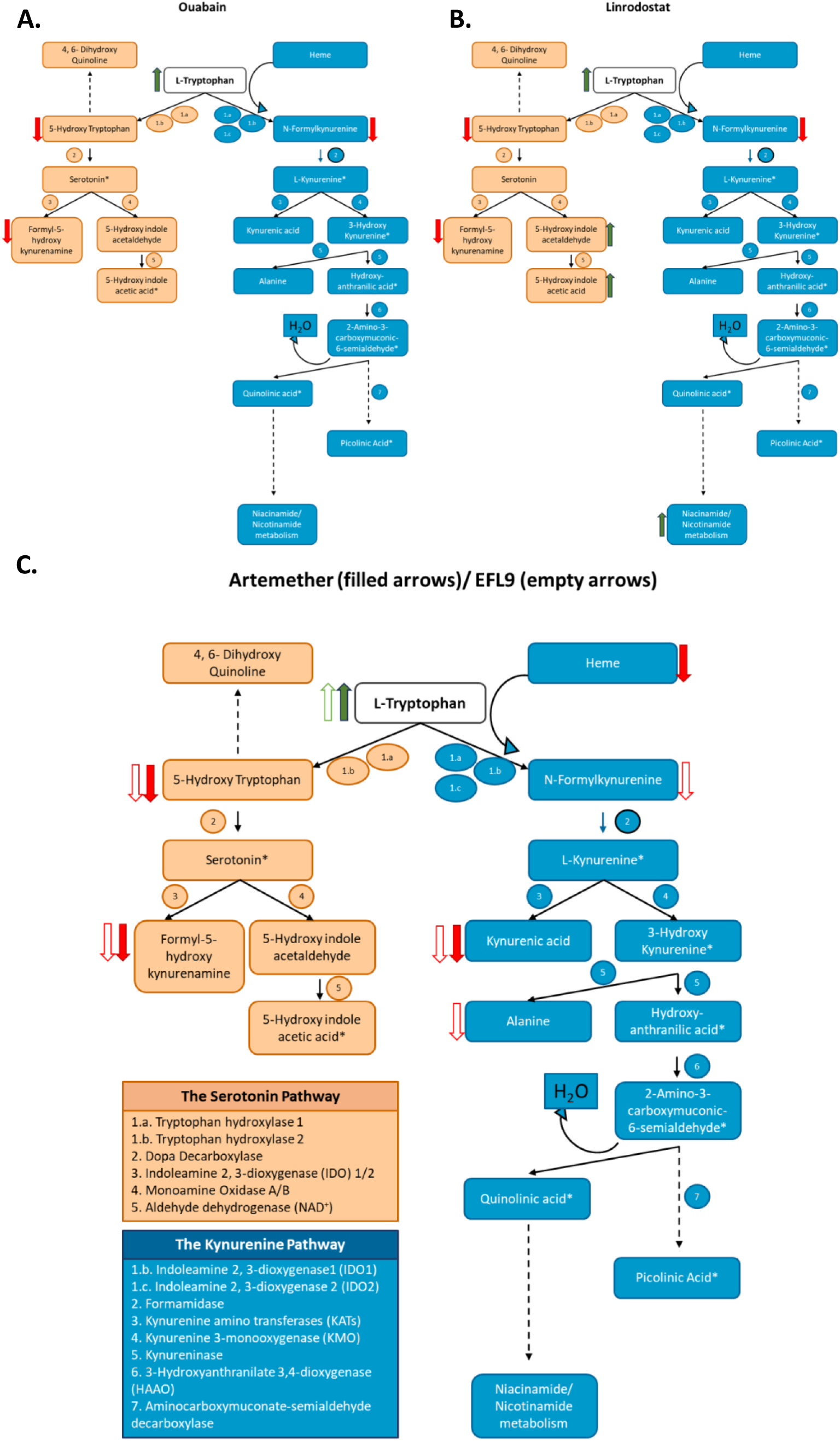
Pathway mapping of intracellular metabolites changing in MDA-MB-231 extracts in response to ouabain, artemether, EFL9 and Linrodostat treatment. **A.** Change in intracellular metabolites upon ouabain treatment in MDA-MB-231 cells. **B.** As in A, but for Linrodostat. **C.** As in A, but for artemether (filled arrows) and EFL9 (empty arrows). Diagrams were made in Microsoft PowerPoint, and maps were created based on Neavin, Liu (92), Stavrum, Heiland (93), Song, Ramprasath (94). Metabolites marked with * were not detectable in our analyses.

Artemether decreased the levels of metabolites downstream of the TPH1/2 initial reaction and decreased kynurenic acid. As mentioned above, artemether was unique in suppressing heme levels. The direct metabolite of the IDO1-mediated conversion of Trp to kynurenine, N-formylkynurenine, was not affected (**Figure 6C**). EFL9, like ouabain, decreased the levels of the immediate reaction product of the IDO1 reaction, N-formylkynurenine, and the downstream metabolites kynurenic acid and L-alanine (a by-product from the synthesis of 3- hydroxyanthranilic acid). EFL9 also affected the serotonin pathway, decreasing levels of the TPH1/2 direct reaction product, 5-hydroxy-L-Trp, as well as decreasing IDO-mediated conversion of serotonin to formyl-5-hydroxykynurenamine **(Figure 6D**).

## Discussion

We investigated the effect of 597 natural products or derivatives on kynurenine production, as an indicator of IDO1 immune checkpoint activity in breast cancer and identified 24 inhibitors and 1 enhancer with low cytotoxic activity. The identification of an IDO1 enhancer might be of interest for conditions were IDO1 is underactive, such as autoimmune diseases (49). Reassuringly, the top inhibitory hit of the screen, alantolactone is a known inhibitor of the JAK/STAT pathway, as well as of other inflammatory mediators including NF-kB (50–52), which are major regulators of IDO1 expression (53–56). Overall, a large subset of the identified hits have been documented to play a role in inflammation, cell cycle progression, PI3K/AKT or MAPK signalling, and several compounds were shown to also have some activity on ion transport. For this study, we decided to focus on two compound classes that showed a clear correlation between chemical structure features and biological activity: artemisinin derivatives and Euphorbia factors. We selected one representative member from each class, artemether and EFL9, and investigated their effect on kynurenine production, the JAK-STAT pathway, and Trp metabolism. We show that both of these drugs are able to downregulate kynurenine levels without interfering with IDO1 expression or STAT1 activation. Rather, artemether suppresses kynurenine production likely through sequestration of heme, by directly binding to the ferrous heme, which can potentially lead to free radical generation, as reported in the literature (45) while EFL9 affects the Trp catabolism pathway in a similar but not identical manner to the IDO1 inhibitor Linrodostat.

Our findings reveal a currently unrecognised role in the regulation of immune metabolism for artemisinin derivatives, which are the WHO recommended treatment for malaria (57–59). Our results enhance our limited understanding of the mechanism of action of artemisinin derived drugs, particularly since we show that the effect of artemether on kynurenine also occurs in non-cancerous cells. The endoperoxide bridge in artemisinin is thought to be the moiety responsible for the compound’s activity. Although the exact target of artemisinin and its mechanism of action are not fully elucidated, one of the popular current hypotheses is that the endoperoxide structure of the drug interacts with Fe(II) released by malaria parasite-induced degradation of haemoglobin, which creates a toxic activated oxygen within the artemisinin molecule (60). Interestingly, artemisinin-related compounds were shown to also have anti-proliferative and pro-apoptotic activity on cancer cells, as well as inhibiting angiogenesis. The exact mechanism of action is unknown; however, increased iron-dependent reactive oxygen species, DNA damage and disrupted mitochondrial membrane have all been proposed as potential mechanistic explanations for these effects (45, 61). Interestingly, in patients, malaria infection with *Plasmodium vivax* was associated with increased kynurenine production and Trp depletion that could not be abolished by treatment with artemether-lumefantrine (62). In our experiments, however, artemether suppressed the kynurenine response in a concentration-dependent manner. Artemether decreased heme levels in breast cancer cells and restoring heme abolished its effect on kynurenine. This is particularly consistent with the structure of the IDO1 enzyme, which is a heme-containing dioxygenase and is believed to exert its mechanism of action by subtracting a proton from the substrate using an iron-bound dioxygen (63). Our findings support that artemether inhibits IDO1 activity by hijacking heme by direct binding from the cytosol and preventing the activation of IDO1. Importantly, this mechanism would impair activity of IDO2 and TDO, which are also metalloenzymes (64), providing an alternative approach to Trp metabolism targeting to the current specific targeting of IDO1.

Of note, artemether also decreased the levels of kynurenic acid, which is an immunoregulatory metabolite. For example, due to its inhibitory effect on Ca^2+^ import, kynurenic acid has been shown to decrease inflammation by decreasing NF-kB activation and to prevent T cell activation and function through decreased cAMP levels (65, 66). Additionally, it has been reported that kynurenic acid inhibits the IL-23/IL-17 pathway (67). Kynurenic acid has also been shown to trigger the dimerization and nuclear translocation of the transcription factor AhR, which subsequently regulates the expression of genes that control the immune response (e.g., increase in IL6 production), but also Trp metabolism (e.g., lack of AhR increases levels of kynurenic acid) (68, 69).

The second class of compounds that we studied included Euphorbia factors. Just as in the case of artemisinin derivatives, all compounds that showed activity had a common chemical feature: the addition of an aromatic moiety on C7 of the core structure. Euphorbia factors have not been investigated in great detail. Some studies have linked the L2 factor with decreased NF-kB-mediated lung inflammation, or decreased hepatocarcinoma progression in mice, associated with STAT3 and Akt inhibition (70, 71). Euphorbia plant derivatives might also have anti-cancer properties, decreasing migration and angiogenetic potential of colon cancer cells, as well as lowering the pool of cancer stem cells (72). Here, we did not observe any effects on STAT1 activation. Euphorbia-derived phytochemicals have also been shown to act as drug resistance modulators, to alleviate HIV-1 and bacterial infections or to disrupt the mitochondrial respiratory chain (73–76). In our study we show that there is a substantial overlap in the metabolic profile of EFL9 and ouabain. This is particularly interesting given the reported effect of some Euphorbia factors on ion transport, which is another common characteristic with the mechanism of action of ouabain (25). However, we identify a specific Trp metabolic signature for EFL9, which is distinct from that of both ouabain and artemether. EFL9 was shown to downregulate both N-formylkynurenine, the direct product of IDO1 activity, and kynurenic acid in cells. A larger number of serotonin pathway metabolites were affected by EFL9, including formyl-5-hydroxykynurenamine, the reaction product of IDO-mediated serotonin degradation decreased in cellular lysates. Similarly, the TPH 1/2 metabolite 5-hydroxy-L-Trp, a serotonin precursor, was also downregulated in cells. EFL9 also downregulated cellular 5- methylthioadenosine, which is an important player in the glutathione pathway, thus maintaining the cellular redox balance (77, 78). Interestingly, EFL9 increased levels of LysPC (lysophosphatidylcholine), 1-18:0-lysoPC and 1-16:0-lysoPC. LysPC is a signalling phospholipid produced by caspase 3-induced Ca^2+^ independent activation of phospholipase A2 (iPLA2), and functions by recruiting phagocytes (79). The elevation in LysPC suggests that EFL9 treatment might thus promote immune mediated tumour-clearance. On the other hand, phosphatidylcholine synthesis, via fatty acid metabolism, is often upregulated in cancer cells, and has been linked to cellular proliferation and cell cycle regulation (80). Thus, further experimental evidence would be required to investigate the anti-cancer and anti-inflammatory effects of EFL9 via phosphatidylcholine/lysophosphatidylcholine upregulation.

Linrodostat is a direct inhibitor of IDO1 which interferes with the heme binding pocket and does not allow the assembly of the enzyme with its iron core (81, 82). Thus, unlike artemether, Linrodostat did not impact the cellular heme levels. Instead, it downregulated the same key metabolites as ouabain: 5-hydroxy-L-Trp, N-formylkynurenine and formyl-5- hydroxykynurenamine, metabolites that were also downregulated by EFL9. However, Linrodostat had a distinct profile from artemether and EFL9, particularly when analysing the list of upregulated metabolites. Linrodostat increased the levels of 5- hydroxyindoleacetaldehyde and indoleacetaldehyde, both of which are metabolites downstream of serotonin, and this indicates potential oxidative stress (83). Linrodostat also upregulated a range of metabolites that were not affected by the other drugs tested in this study. The unique metabolic signature of Linrodostat suggests that neither artemether nor EFL9 fit the profile of a direct IDO1 inhibitor. However, we report overlapping metabolic profiles for EFL9, ouabain and Linrodostat in cells, which will need to be further studied in future biochemical and functional studies.

In conclusion, by screening a library of 597 NPs and their derivatives, predominantly terpenoids, for effects on kynurenine production by triple-negative breast cancer cells, we discovered two inhibitors, artemether and EFL9. We identify chemical features that are required for the biological activity of these compounds. By studying the Trp catabolic pathway, and by comparison with the previously described IDO1 regulator ouabain (25) and with a clinically used selective IDO1 inhibitor (Linrodostat), we demonstrate distinct metabolic profiles, with ouabain and EFL9 showing the largest overlap in terms of key Trp metabolites. We show that artemether suppresses the kynurenine response by interfering with intracellular heme availability, a mechanism that would affect all members of the IDO/TDO family of metalloenzymes and could be relevant to its clinical use as an anti-malarial drug. Importantly, we demonstrate that the kynurenine-suppressing activity of artemether and EFL9 is not specific to cancer cells. Overall, our findings set the foundation for future studies to explore these NPs and/or their chemically modified derivatives as novel tools to regulate Trp metabolism in cancer and other pathologies.

## Methods

### Cell culture

#### Culture conditions

Human triple-negative metastatic breast cancer MDA-MB-231 cells were provided by our collaborator, Mustafa Djamgoz. Human primary mammary epithelial (HMEC CC-2551) cells from 2 donors were purchased from Lonza. All experiments with MDA-MB-231 and HMEC cells were conducted at 37 °C, 5% CO_2_. MDA-MB-231s were cultured in Dulbecco’s Modified Eagle’s Medium (DMEM) (21969035, Gibco) supplemented with 10 % heat-inactivated Foetal Calf Serum (FCS), 1 % L-glutamine (25030081, Gibco), and 1 % penicillin/ streptomycin (15070063, Gibco). Human mammary epithelial cells (HMECs) were cultured using Mammary Epithelial Cell Growth Medium Bullet Kit (MEGM) (CC-3150, Lonza), passaging was carried out using the Reagent Pack Subculture Reagents for primary cell sub culturing (CC-5034, Lonza).

### Seeding densities

Drug titrations were carried out in MDA-MB-231 cells, seeded at 2.5 x 10^5^/ well in 1 mL complete DMEM in 12-well plates. Metabolomics analysis was carried out in MDA-MB-231 cells seeded at 4x10^6^/ dish in 10 cm dishes. HMECs were seeded at 0.5 x 10^5^/ well in 100 µL media in clear 96-well flat-bottom plates.

### Kynurenine assay

For each kynurenine assay a set of 8 standards, ranging from 0 to 100 μM, was prepared, using a 50 mM in 0.5 M HCl stock. Then, 140 µL cell culture supernatants were transferred to a U-bottom 96-well plate, alongside the standards. Wells were treated with 10 µL 30 % 6.1 N trichloroacetic acid for 30 min at 50 °C. On hundred µL of each sample/ standard were then transferred to a new 96-well flat-bottom plate and topped up with 100 µL 1.2 % w/v p-dimethylaminobenzaldehyde in glacial acetic acid. Following a 10-min RT incubation, absorbance at 492 nm was measured on a VersaMax microplate reader (Molecular Devices), using the SoftMax Pro data acquisition software (40, 41).

### Viability

Viability was mainly assessed through automatic cell counting. Cells were lifted with 0.05 % Trypsin-EDTA (25300054, Gibco), stained with trypan blue (T10282, Themo Fisher Scientific), and counting was done automatically using a Countess II FL Automated Counter (C10283, Thermo Fisher Scientific).

A plate-based measurement was used to assess viability for experiments where counting individual cells was considered detrimental, i.e. due to large number of samples, or to reduce cell exposure to stress (e.g., the compound screen, for primary HMECs experiment). For plate-based measurements, the Deep Blue Viability dye was added to the cell cultures at a 1:10 ratio with the supernatant volume. Cells were incubated at 37 °C, 5 % CO_2_ for 4 h and viability was then measured by reading optical density (OD) at 570 and 600 nm, using a VersaMax (Molecular Devices) plate reader. The absorbance value recorded at 600 nm was then subtracted from the 570 nm absorbance value, to give a measure of cell viability. The given numbers were then normalized (fold change calculations) to the corresponding control (the most viable condition, according to the experimental design) to give a ratio corresponding to sample viability.

### Natural product screen

#### Set-up

The drug screen was carried out using MDA-MB-231 cells, seeded at 7x10^4^ cells/ well in 100 μL fully supplemented DMEM, in 96-well tissue culture treated plates, 24 h prior drug treatment.

Six hundred thirty conditions were screened, corresponding to 597 unique compounds as some NPs were included both in the MCE and in-house libraries. They were split in 10 plates, each with a similar set-up, as in **Supplementary Figure S2**. All plates had 3 unstimulated controls, 3 positive controls consisting of cells treated with IFNγ/ TNF and 3 negative controls of cells treated with IFNγ/ TNF and 200 nM ouabain. Drug plates were all prepared at the same time, by aliquoting 1 μL per well of 10 mM drug solution in 100 % DMSO in 700 μL 96- well v-bottom plates. For ouabain controls, 4 μL of 50 μM ouabain 1 % DMSO topped up with 0.96 μL 100 % DMSO were added to corresponding wells. For DMSO controls 1 μL DMSO per well was added. The plates were then stored at -80 ^◦^C until the day of the screen. On this day, compounds were topped up with complete DMEM and added to the cells to give a final well volume of 250 μL, with a final well concentration for natural compounds of 10 μM in 0.1 % DMSO (**Supplementary Figure S6**). The well concentration for ouabain was 200 nM, in 0.1 % (v/v) DMSO, as selected based on previous experiments (25). Each drug plate was tested in duplicate. For each plate of compounds, the duration of the screen was 4 days. Drug pre-treatment was carried out for 18 h. Cytokine stimulation was then added straight into the culture media for another 24 h at final well concentration of 1 U/ mL for IFNγ and 6.25 ng/ mL for TNF to induce expression of IDO1 (84). Supernatants were collected for the kynurenine assay and remaining cells were used to measure viability using the Deep Blue viability assay. *Analysis. Selection of Compounds Of Interest (COIs) and hits*.

Two selection criteria based on the effect of compounds on kynurenine were used for the screen. The standard deviation (SD) variation method was applied by firstly calculating the SD for the vehicle control. Next the following formula was applied to calculate variation:

X = (Sample Kyn – Mean Vehicle Kyn)/Vehicle SD

The fold change method used the following formula:

Y = Sample Kyn/Mean Vehicle Kyn

In addition to the kynurenine-based selection criteria, the following formula was used to calculate viability fold change:

Z = Sample Viability/Mena Vehicle Unstimulated viability.

For the first round of screening compounds that gave a fold change in kynurenine levels ≤ 0.5 or ≥ 1.4 OR a change in kynurenine equivalent to ≥ 3 or ≤ -3 SDs away from the vehicle control, while maintaining viability fold change ≥ 0.7, were considered COIs. Both replicate plates were included in the fold change and viability analyses, while for the SD variation analysis only plates where the controls were separated by an SD variation of at least 3 were included, replicate plates that did not pass this quality control were used to qualitatively verify patterns for potential COI exclusion. For the validation round hits had to cause both a fold change in kynurenine levels ≤ 0.5 or ≥ 1.4 and a change in kynurenine equivalent to ≥ 3 or ≤ -3 SDs away from the vehicle control while maintaining viability fold change ≥ 0.7.

### Drug and NP treatment assays

Chemical formulas were taken from the supplier website (ChemFaces) or the publicly available database PubChem (PubChem CID: 42608244). For drug titrations, MDA-MB-231 cells were pre-treated with drugs at 0.1, 1 and 10 µM concentrations in 0.1% DMSO, or 200 nM in 0.1% DMSO for ouabain for 18 h, then the culture supernatants were replaced with fresh drug solution supplemented with IFNγ (1 U/ mL)/ TNF (6.25 ng/ mL) for another 24 h. Supernatants were then collected for the kynurenine assay.

For the metabolomics experiments, MDA-MB-231s were treated with 200 nM ouabain 0.1% DMSO, or 10 µM of the corresponding drug in 0.1% DMSO for 18 h, and the cytokines were supplemented directly into the culture media for another 24 h, to a final concentration of IFNγ (1 U/ mL)/ TNF (6.25 ng/ mL). Kynurenine assays were performed on the supernatants. Supernatant and cell lysates were collected using the freeze-thaw protocol described in Rushing, Schroder and Sumner (85). For further mechanistic experiments, the heme scavenger succinyl acetone from Cambridge Bioscience LTD (25501-10mg-CAY) and the heme stabiliser hemin (porcine) from VWR International (A11165.06) were used.

For primary cell (HMEC) experiments, drug treatment was carried out with 200 nM ouabain or 10 µM corresponding drug in 0.1% DMSO for 18 h. Cytokines were then supplemented to the culture medium to give a final concentration of IFNγ (100 U/ mL)/ TNF (6.25 ng/ mL). Twenty-four h post cytokine stimulation kynurenine assays were performed on supernatants and viability was measured using the Deep Blue viability assay.

### RNA extraction and qRT-PCR

QIAzol (79306, Qiagen) and RNeasy Mini kit (74104, Qiagen) were used to extract total cellular mRNA. cDNA synthesis was performed using random hexamers and Superscript II (Invitrogen) reverse transcriptase (18064014, Invitrogen). qRT-PCR of IDO1, PD-L1, GAPDH and actin was performed using Fast SYBR Green qRT-PCR (4385610, Applied Biosystems). Human IDO1 was quantified using a QuantiTect Primer Assay (Qiagen). Human PD-L1 and β-actin were quantified using the following forward and reverse primers (Sigma Aldrich): PD- L1 Forward 5′-CATCTTATTATGCCTTGGTGTAGCA-3′, PD-L1 Reverse 5′- GGATTACGTCTCCTCCAAATGTG-3′, β-actin Forward: 5-CACCATTGGCAATGAGCGGTTC-3’, β-actin Reverse 5’-AGGTCTTTGCGGATGTCCACG-3’. Primers were used at a working concentration of 10 µM. Relative mRNA levels were calculated using the 2^-ΔCT^ method. mRNA levels were normalized to β-actin. The 2^-ΔCT^ value was calculated for each sample after β-actin normalisation and fold change from the average 2^-ΔCT^ value across all the samples was calculated for each condition.

### Western blots

Cold radioimmunoprecipitation assay (RIPA) lysis buffer (5 mM EDTA, 150 mM NaCl, 10 nM Tris-HCl, pH 7.2, 0.1% SDS, 0.1% Triton X-100, and 1% sodium deoxycholate) supplemented with protease inhibitory complexes P8340, P5726, and P0044 (Sigma) (1 in 100 dilution) was used for harvesting and lysing the cells. Protein concentration was determined using a Pierce bicinchoninic acid assay (23225, Thermo Scientific). For sample electrophoresis 10% SDS-PAGE gels were used and transfer was carried out onto PVDF membranes (IPVH00010, Millipore). Two % BSA was used for the blocking step for a minimum of 15 min at room temperature. The following antibodies were probed overnight at 4 °C (1:1000): anti-PD-L1 (E1L3N), anti-STAT1 (9172), anti-pSTAT1 Tyr-701 (D4A7), anti-IDO1 (D5J4E) all from Cell Signaling Technology. Antibodies for GAPDH (6C5) from Abcam were probed for 1 h at room temperature. Membranes were further incubated with horseradish peroxidase (HRP)- conjugated secondary antibodies (P044701-2 or P044801-2, Dako Agilent) and visualized with ECL (10754557, GE Healthcare Amersham) on film (28906836, GE Healthcare) and developed on a Medical Film Processor SRX-101A (Konica Minolta). Band intensity was quantified using Fiji 2.0.0-rc-69/1.52p (NIH, Bethesda, MD) (96).

### Targeted Trp catabolism metabolomics

#### Set-up

For metabolomics experiments, cells were seeded at 4x10^6^/ dish in 10 cm dishes, and treated with 200 nM ouabain 0.1 % DMSO, or 100 nm Linrodostat (Generon, M6248) in 0.1% DMSO or 10 µM of the corresponding natural compound (artemether and Euphorbia factor L9, ChemFaces, CFN99184 and CFN92887, respectively) in 0.1 % DMSO for 18 h. A DMSO-only control was also set up. Cytokines were supplemented directly into the culture media for another 24 h to a final concentration of IFNγ (1 U/mL)/TNF (6.25 ng/mL). Kynurenine assays were performed on the supernatants. Supernatant and cell lysates were collected using the freeze-thaw protocol described in Rushing, Schroder and Sumner (85).

Two µL of sample were injected onto a Thermo Vanquish Flex Liquid Chromatography (LC) system fitted with a Waters High Strength Silica T3 column (100 x 2.1 mm). Mobile phases were: A, water + 0.1% formic acid (v/c); B, methanol + 0.1% formic acid (v/v). Samples were eluted over a 10 min gradient as follows: 0-16 min 2-100 % B, 16-20 min isocratic 100 % B, 20-20.1 min to 2 % B, 20.1-30 min re-equilibrate at 2 % B. The column was held at 50 °C and flow rate was 0.4 mL/min. The eluate was sent to a Thermo Orbitrap Fusion instrument between 0.6 and 28 min, fitted with a Heated Electrospray Ionization source operating in positive ion mode (sheath gas 50 units, auxiliary gas 10 units, sweep gas 1 unit, spray voltage 3500V, vaporizer 325°C, ion transfer tube 275°C). The mass spectra (MS) 1 and MS2 data were completed at an orbitrap resolution of 60,000 full width at half maximum (FWHM), with a 0.4 s cycle time. MS1 data was collected between 70 - 1050 m/z. Easy internal calibration was used to achieve mass accuracy < 1 parts per million (ppm). MS2 data was collected alternating between Higher Collisional Dissociation mode with stepped collision energies (35, 40, 45 %) and Collision-Induced Dissociation Mass Spectrometry mode with a fixed collision energy of 40 %. Dynamic exclusion was set for n = 1 over 4s. Once collected, all files were converted to centroided. mzML and mgf formats using ProteoWIzard msconvert.

### Analysis

For metabolomics analysis data were processed through bespoke XCMS scripts (provided by the Bioscience Technology Facility at the University of York) to look for exact MS1 matches of the Trp pathway metabolites, with annotations verified and additional annotations included using high-resolution MS2 data processed through Sirius.

The raw data files were converted to centroided mzML and mgf formats using ProteoWIzard msconvert. The mzML format files were used as inputs for feature detection using bespoke scripts written in R with functions from the XCMS (86–88). Features were detected with the centWaveWithPredictedIsotopeROIs method, with the following function parameters: ppm = 10; signal to noise threshold (snthresh) = 3; range of peak elution times (s) (peakwidth) = c (2, 10); peak definition: number of data points exceeding a set intensity threshold (prefilter) = c (3, 1000); integration method for peak quantification (integrate) = 2 (real data); accepted closeness of two signals in m/z dimension (mzdiff) = -0.1. Duplicate features and features outside the time between 0.7 and 28 min were removed using custom scripts. Features were subsequently grouped using a modified *scms group()* function. Adducts and isotopes were detected using CAMERA. Feature MS1 values were matched to a custom database comprising predicted [M+H] + or [M+H] + ions of the Trp pathway as defined by pathway ID SMP0000063 in the HMDB (89). The best representative MS2 spectra (defined as nearest to the MS1 precursor feature apex) were extracted with custom scripts from the .mgf files and converted to .ms2 format using custom scripts for input into Sirius. Feature tables were initially filtered to: 1) only retain features found at least once in any sample with an area value greater than the mean + 3 x the SD value of blank samples; 2) retain any features with a HMDB database hit; and 3) retain features that are probably mono-isotopes (as calculated by CAMERA). Sirius matching was then performed on available initially filtered features, using Sirius 5.8.2 (90, 91) all default databases and instruments were set to Orbitrap with 5 ppm tolerance. Sirius outputs were merged with feature tables. These were manually further edited to remove redundant and near-baseline features and reconcile HMDB and Sirius annotated features.

### Materials and natural products

403 NPs were purchased from MedChemExpress. The rest were generated in-house or purchased from Merck, Insight Biotechnology, ChemFaces, SLS, Generon, LC labs, Muse Chem, TRC, or VladaChem (**Supplementary Figure S1**). Deep Blue Cell Viability Kit (424702) was purchased from BioLegend. Human recombinant IFNγ (300–02) was purchased from PeproTech and TNF (210-TA-020) was obtained from Biotechne R&D systems. Ouabain octahydrate (O3125), (L-kynurenine (K8625), p-dimethylaminobenzaldehyde (D2004), were purchased from Sigma Aldrich. Trichloroacetic acid (11462691) and glacial acetic acid (010994-AC) were purchased from Fisher Scientific. Cell culture grade DMSO (A3672.0100) was purchased from AppliChem. Trypan blue (T10282) was purchased together with Countess™ Cell Counting Chamber Slides (C10283) from Thermo Fisher Scientific. Methanol was purchased from Honeywell (32213–2).

### Statistical analysis

For all data (except for the metabolomics analysis) statistical analysis was carried out in Graph Pad Prism v9.0.0. All experiments were carried out with up to 4 biological repeats, therefore, the sample size was too small to perform normality analysis confidently, and normal distribution was assumed for all data. All data were thus plotted as means and SD, and statistical analysis was performed using an ordinary one-way ANOVA, coupled with a Bonferroni’s post-test (when specific pairs of conditions were compared), or a Tukey’s post-test when all conditions were compared against each other.

For statistical analysis of the metabolomics data, the Metaboanalyst software was used. Metaboanalyst applied a data integrity check to each set of data uploaded, excluding metabolite entries that exhibited constant or no value for all the treatment conditions included in the data set. All data were median normalised and log10 transformed. Principal component analysis was used to verify correct clustering of the data based on treatment conditions. Further statistical analysis was carried out on annotated data only. The data were split in 3 files containing paired conditions. A data integrity check was applied to each set of data resulting in exclusion of metabolite entries that exhibited constant or no value for all the treatment conditions included in the data set. Comparisons were carried out using an unpaired t-test and fold change analysis. Differentially abundant metabolites of were identified based on the following thresholds: FDR ≤ 0.1 and fold change value ≥ 1.5.

## Data availability statement

The data included in this study (MTBLS10694) are in the ’In Curation’ stage and will be available here: https://www.ebi.ac.uk/metabolights/MTBLS10694. All other raw data are available on request.

## Acknowledgments

We thank Tony Larson and Adam Dowle at the Metabolomics and Proteomics Lab in the University of York Bioscience Technology Facility for assistance with Trp metabolomics. The authors also thanks Leonardo Gomez and Rachael Simister for providing the supercritical CO_2_ fucoxanthin extract, and Carla Machado for the fucoidans extracted from pelagic sargassum.

## Author contributions

A.L.C.: Data Curation, Investigation, Formal Analysis, Methodology, Validation, Writing Original Draft, Writing Review and Editing.

T.C.: Data (NP Library) Curation, Formal Analysis, Funding acquisition, Writing Review and Editing.

C.P-J.: Data Curation, Investigation, Formal Analysis, Validation, Writing Review and Editing.

T.T.: Data (NP Library) Curation, Formal Analysis, Funding acquisition, Writing Review and Editing.

I.K.: Data Curation, Formal Analysis, Funding acquisition, Writing Review and Editing.

B.R.L.: Data (NP Library) Curation, Formal Analysis, Funding acquisition, Writing Review and Editing.

W.J.B.: Conceptualisation, Funding acquisition, Supervision, Writing Review and Editing. I.A.G.: Conceptualisation, Funding acquisition, Supervision, Writing Review and Editing.

D.L.: Conceptualisation, Funding acquisition, Supervision, Project Administration, Formal Analysis, Writing Original Draft, Writing Review and Editing.

## Funding and additional information

ALC was funded by the UKRI BBSRC White Rose doctoral training partnership (BB/J014443/1). Additional support was received from a UKRI BBSRC Food Agriculture and Health Project Award managed by the University of York (BB/W510737/1) and from a UKRI MRC Impact Accelerator Award (MR/X502662/1) also managed by the University of York.

