## Supplementary Figures for "Artemether and Euphorbia Factor L9 suppress kynurenine production through distinct effects on Tryptophan metabolism"

**Supplementary Figures S1-S6**

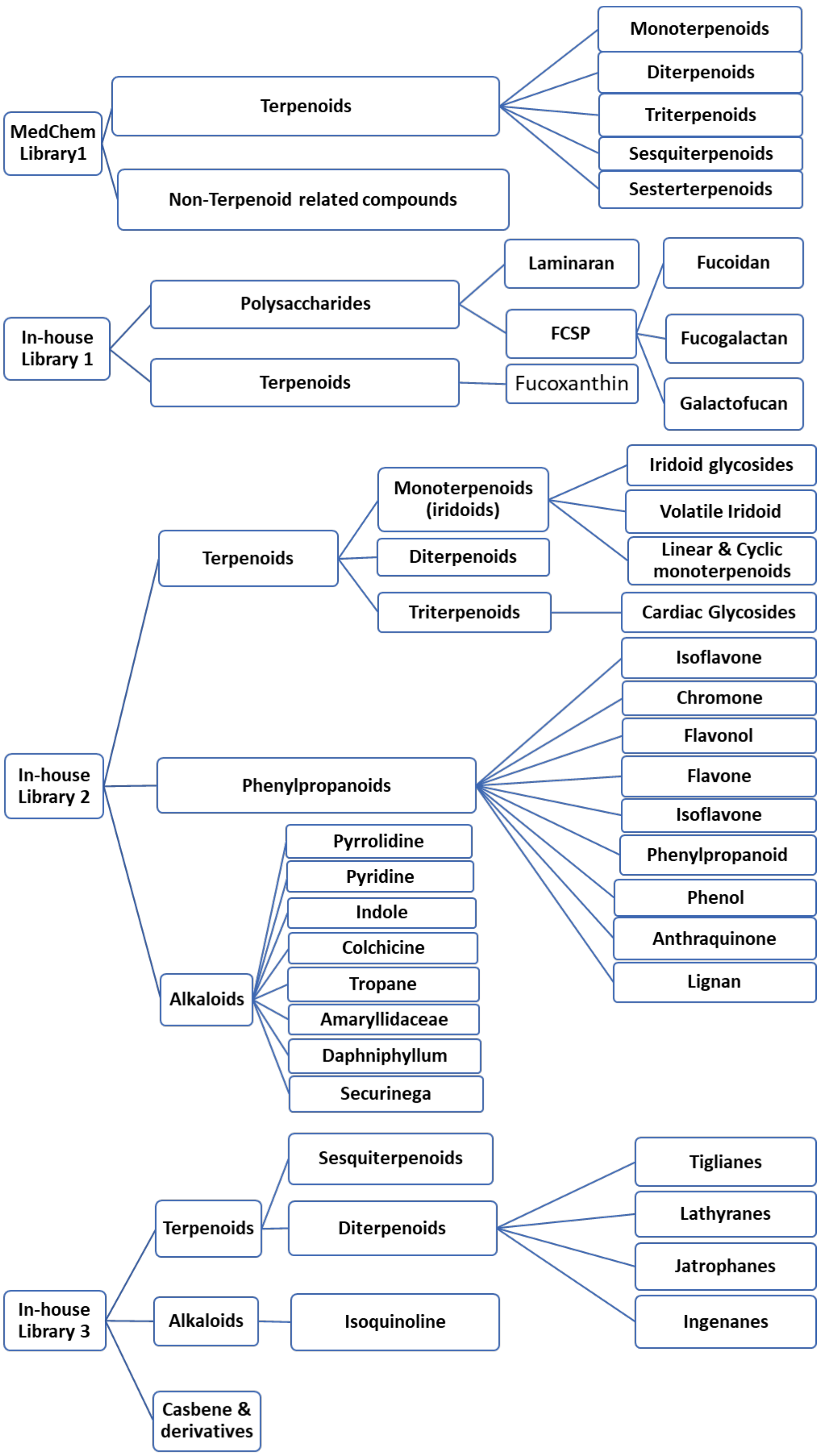


**Supplementary Figure S1.** Summary of the main classes of 597 compounds and their sources: commercially available MedChemExpress (MCE) library and three in-house generated libraries. FCSP - Fucose-containing sulphated polysaccharides.

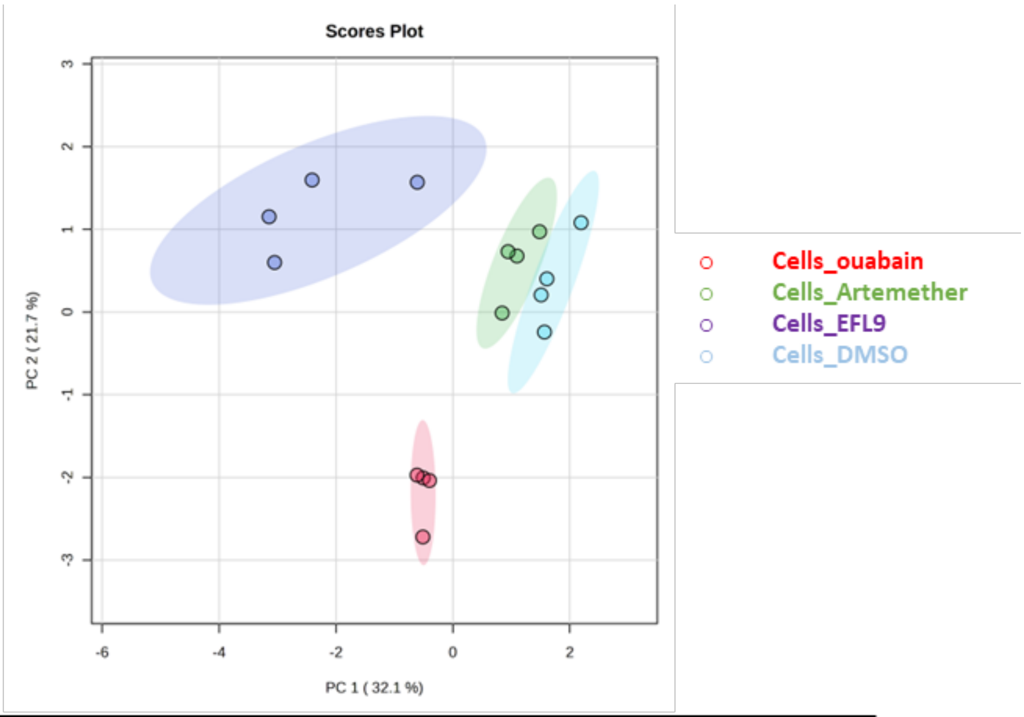


| PC1 | PC2 | PC3 | PC4 | PC5 |
|---|---|---|---|---|
| 32.1% | 21.7% | 12.1% | 9.9% | 6.2% |

**Supplementary Figure S2.** Impact of ouabain, artemether and EFL9 on Trp metabolites in MDA-MB-231 cell extracts. PCA analysis of the entire metabolite output data set (135 metabolites) in cells treated with DMSO (blue), artemether (green), EFL9 (purple), and ouabain (red), n=4 biological replicates.

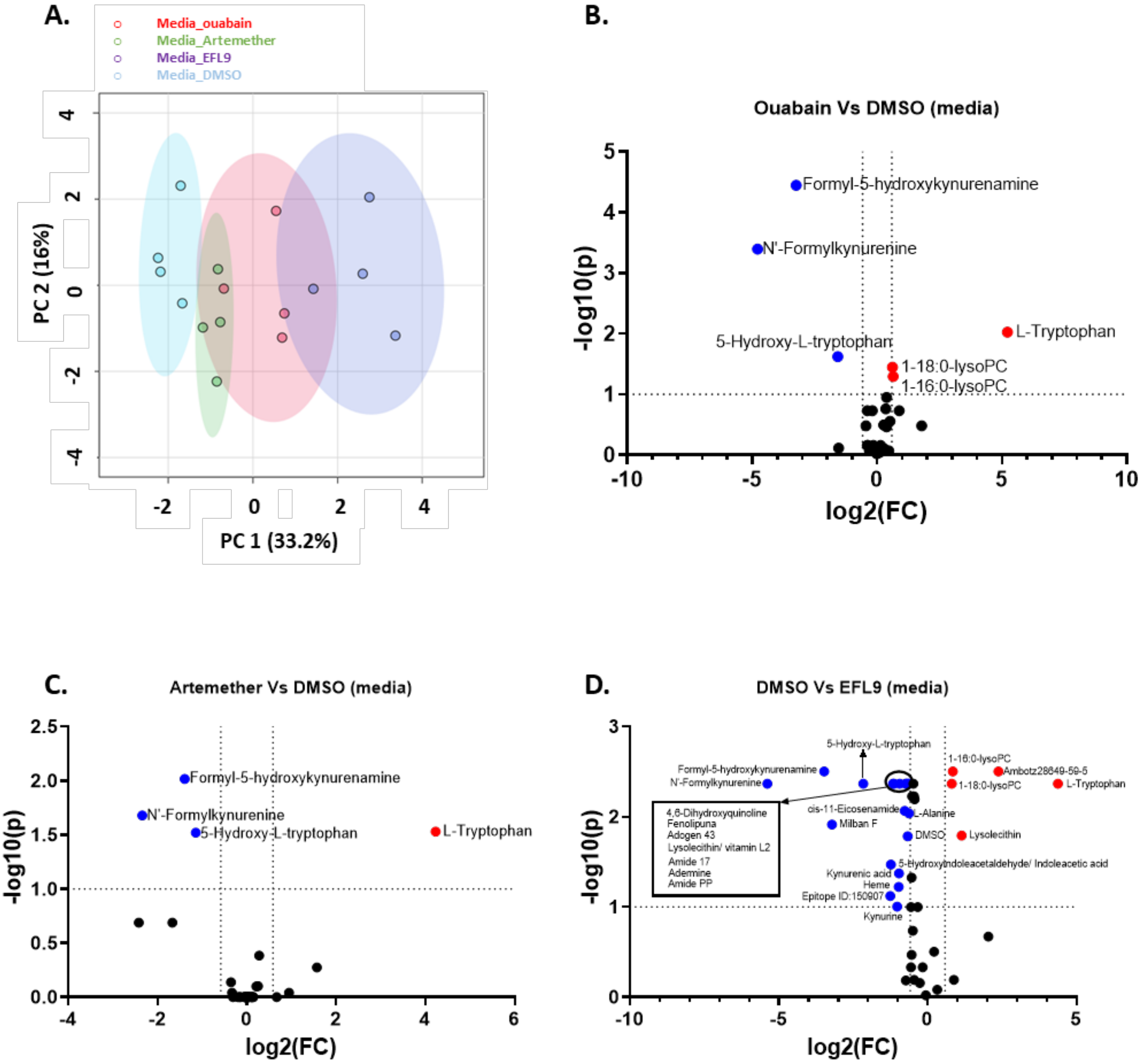


**Supplementary Figure S3.** Impact of ouabain, artemether and EFL9 on Trp metabolites secreted by MDA-MB-231 cells. A. PCA analysis of the annotated metabolite data set (135 metabolites with 19 metaboanalyst automatic exclusions), n=4. B. Volcano plot of log10(Fold change) Vs log10(p) looking at changes in metabolites in response ouabain, analysis was carried out using the annotated metabolite data set (47 metabolites with 6 metaboanalyst automatic exclusions), n=4. C. Volcano plot of log10(Fold change) Vs log10(p) looking at changes in metabolites in response artemether, analysis was carried out using the annotated metabolite data set (47 metabolites with 5 metaboanalyst automatic exclusions), n=4. D. Volcano plot of log10(Fold change) Vs log10(p) looking at changes in metabolites in response EFL9, analysis was carried out using the annotated metabolite data set (47 metabolites with 4 metaboanalyst automatic exclusions), n=4.

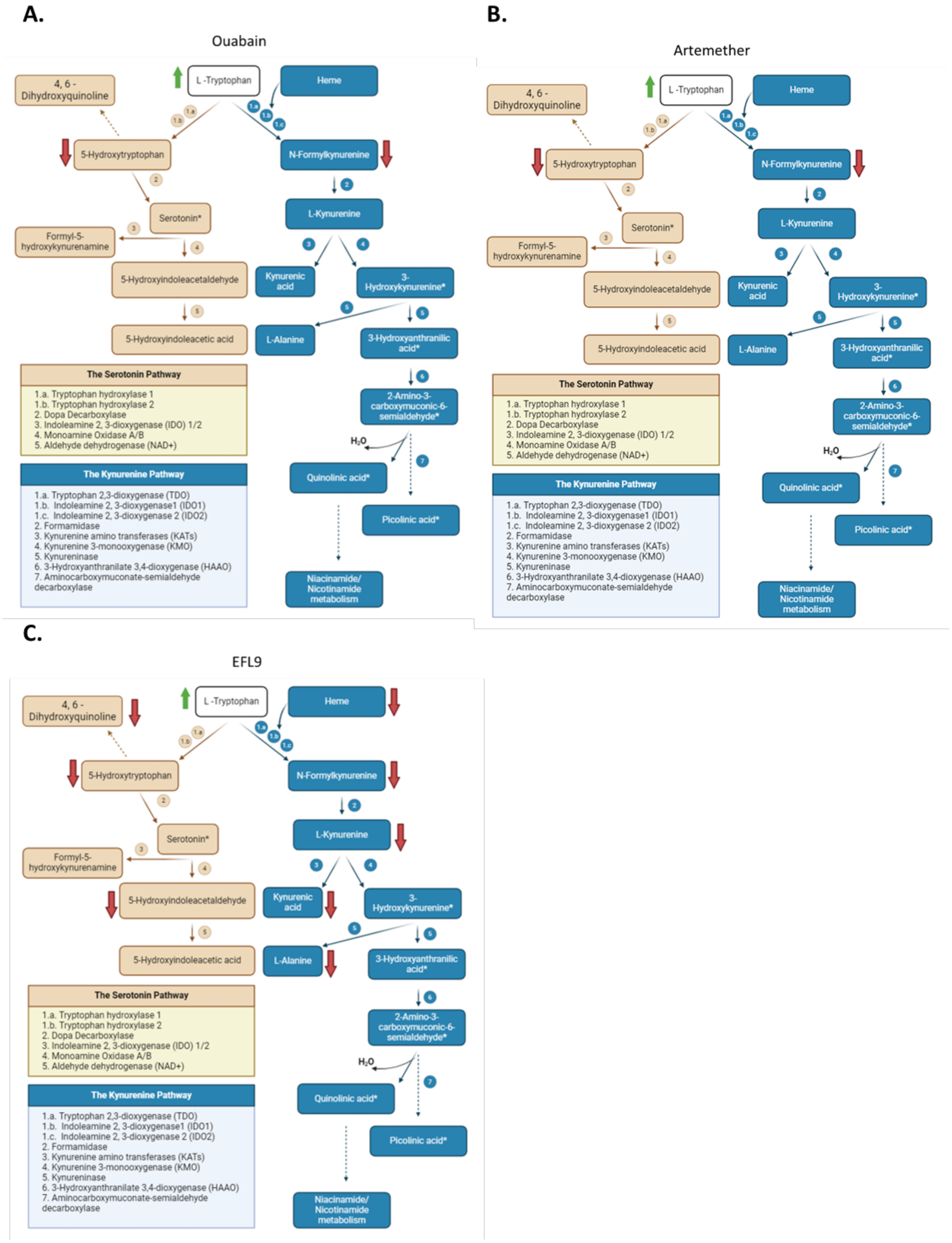


**Supplementary Figure S4.** Summary of the impact of ouabain (A), artemether (B) and EFL9 (C) on secreted Trp metabolites in MDA MB 231 cultures. Red arrows indicate a decrease, green arrow indicate an increase.

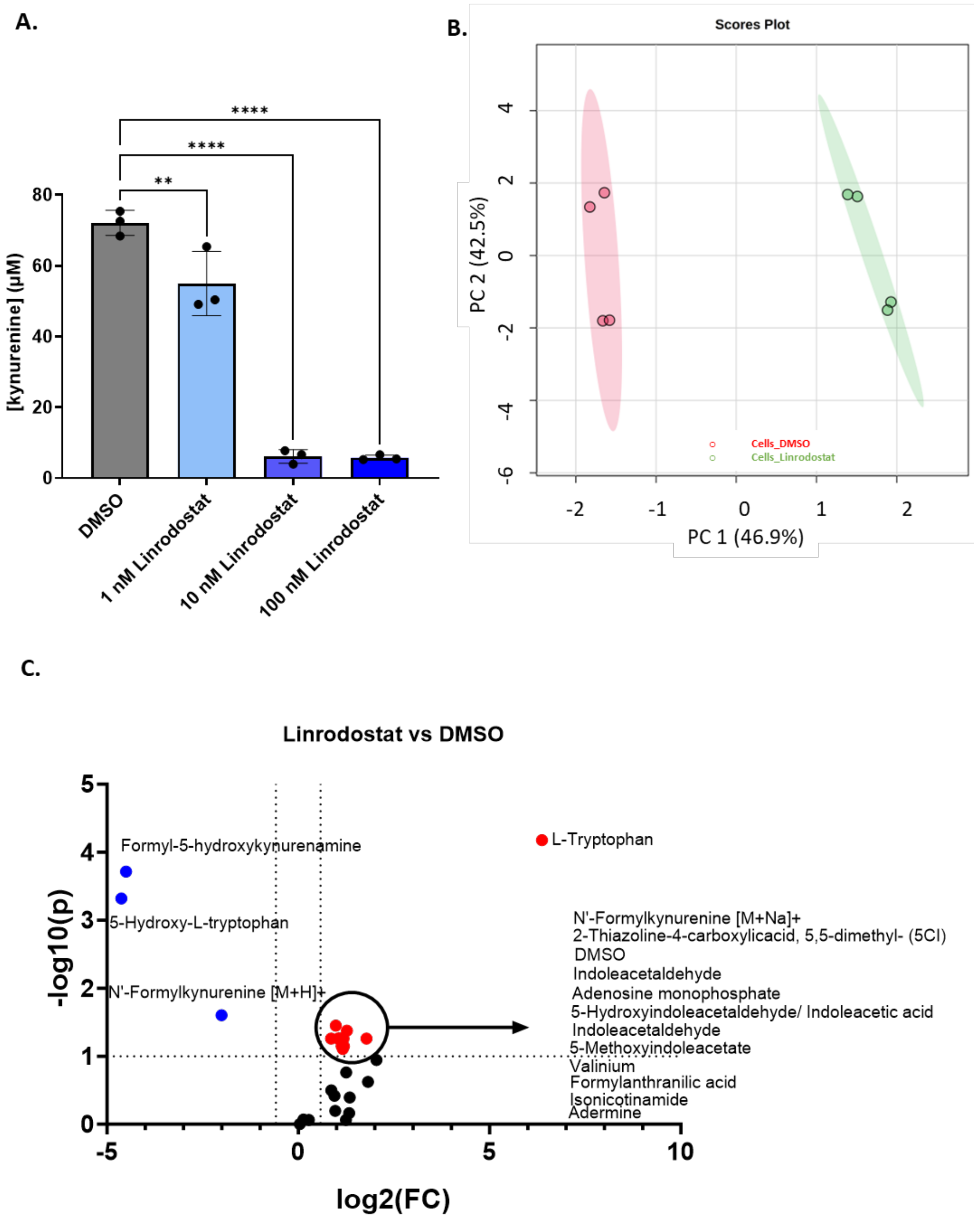


**Supplementary Figure S5.** Impact of Linrodostat on Trp metabolism in MDA-MB-231 cells. A. Effect of the commercially available IDO1 inhibitor, Linrodostat, on kynurenine production in MDA-MB-231 cells, n=3 biological replicates. Statistical analysis was carried out using a one-way ANOVA coupled with a Bonferroni's post-test (all conditions were compared to the DMSO control), for specific comparisons ( **p<0.01; ****p<0.0001) B. PCA analysis of the annotated metabolite data set (28 metabolites, n=4). C. Volcano plot of log10(Fold change) Vs log10(p) looking at changes in metabolites in response Linrodostat, analysis was carried out using the annotated metabolite data set (28 metabolites), n=4.

**A.**

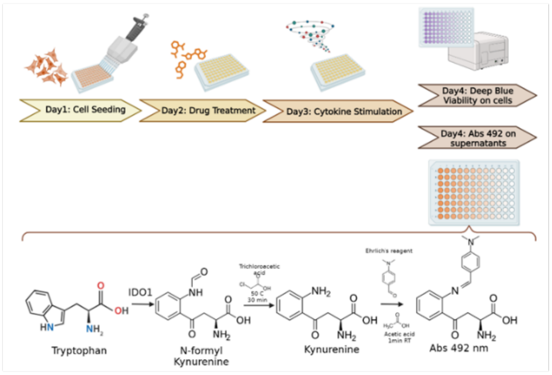


**B.** **Drug plate**

| | 1 | 2 | 3 | 4 | 5 | 6 | 7 | 8 | 9 | 10 | 11 | 12 |
|---|---|---|---|---|---|---|---|---|---|---|---|---|
| A | | | | vehicle unstimulated | 1 | 2 | 3 | vehicle | 4 | 5 | 6 | Ouabain |
| B | | | | 7 | 8 | 9 | 10 | 11 | 12 | 13 | 14 | 15 |
| C | | | | 16 | 17 | 18 | 19 | 20 | 21 | 22 | 23 | 24 |
| D | | | | Ouabain | 25 | 26 | 27 | vehicle unstimulated | 28 | 29 | 30 | vehicle |
| E | | | | 31 | 32 | 33 | 34 | 35 | 36 | 37 | 38 | 39 |
| F | | | | 40 | 41 | 42 | 43 | 44 | 45 | 46 | 47 | 48 |
| G | | | | 49 | 50 | 51 | 52 | 53 | 54 | 55 | 56 | 57 |
| H | | | | vehicle | 58 | 59 | 60 | Ouabain | 61 | 62 | 63 | vehicle unstimulated |

| | 1 | 2 | 3 | 4 | 5 | 6 | 7 | 8 | 9 | 10 | 11 | 12 |
|---|---|---|---|---|---|---|---|---|---|---|---|---|
| A | 100 | 100 | 100 | vehicle unstimulated | 1 | 2 | 3 | vehicle | 4 | 5 | 6 | Ouabain |
| B | 50 | 50 | 50 | 7 | 8 | 9 | 10 | 11 | 12 | 13 | 14 | 15 |
| C | 25 | 25 | 25 | 16 | 17 | 18 | 19 | 20 | 21 | 22 | 23 | 24 |
| D | 12,5 | 12,5 | 12,5 | Ouabain | 25 | 26 | 27 | vehicle unstimulated | 28 | 29 | 30 | vehicle |
| E | 6,25 | 6,25 | 6,25 | 31 | 32 | 33 | 34 | 35 | 36 | 37 | 38 | 39 |
| F | 3,13 | 3,13 | 3,13 | 40 | 41 | 42 | 43 | 44 | 45 | 46 | 47 | 48 |
| G | 1,56 | 1,56 | 1,56 | 49 | 50 | 51 | 52 | 53 | 54 | 55 | 56 | 57 |
| H | 0 | 0 | 0 | vehicle | 58 | 59 | 60 | Ouabain | 61 | 62 | 63 | vehicle unstimulated |
| | Kynurenine standards | | | | | | | | | | | |


| | 1 | 2 | 3 | 4 | 5 | 6 | 7 | 8 | 9 | 10 | 11 | 12 |
|---|---|---|---|---|---|---|---|---|---|---|---|---|
| A | 100 | 100 | 100 | vehicle unstimulated | 1 | 2 | 3 | vehicle | 4 | 5 | 6 | Ouabain |
| B | 50 | 50 | 50 | 7 | 8 | 9 | 10 | 11 | 12 | 13 | 14 | 15 |
| C | 25 | 25 | 25 | 16 | 17 | 18 | 19 | 20 | 21 | 22 | 23 | 24 |
| D | 12,5 | 12,5 | 12,5 | Ouabain | 25 | 26 | 27 | vehicle unstimulated | 28 | 29 | 30 | vehicle |
| E | 6,25 | 6,25 | 6,25 | 31 | 32 | 33 | 34 | 35 | 36 | 37 | 38 | 39 |
| F | 3,13 | 3,13 | 3,13 | 40 | 41 | 42 | 43 | 44 | 45 | 46 | 47 | 48 |
| G | 1,56 | 1,56 | 1,56 | 49 | 50 | 51 | 52 | 53 | 54 | 55 | 56 | 57 |
| H | 0 | 0 | 0 | vehicle | 58 | 59 | 60 | Ouabain | 61 | 62 | 63 | vehicle unstimulated |
| | Kynurenine standards | | | | | | | | | | | |


**C.**

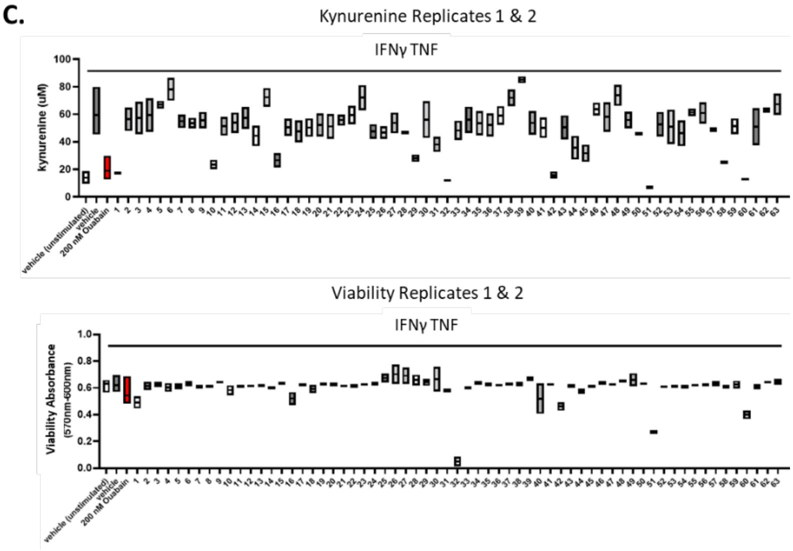


**Supplementary Figure S6.** Drug screen set-up and representative raw data per plate. A. Drug screen workflow: the kynurenine assay and deep blue viability; B. Drug plate and kynurenine plate maps for each set of compounds; C. Representative raw data for the first set of compounds.

### Supplementary Tables

#### Supplementary Table S1: Summary of drug screen results

Supplementary Table1.xlsx

#### Supplementary Table S2: Drug hits and their biological activity

Supplementary Table 2.xlsx
